# Theta and alpha oscillations in human hippocampus and medial parietal cortex support the formation of location-based representations

**DOI:** 10.1101/2023.10.18.562859

**Authors:** Akul Satish, Vanessa G. Keller, Sumaiyah Raza, Shona Fitzpatrick, Aidan J. Horner

**Affiliations:** Department of Psychology, University of York, UK; MRC Cognition and Brain Sciences Unit, University of Cambridge, UK; York Biomedical Research Institute, University of York, UK

**Keywords:** Spatial Memory, Scene Integration, Associative Memory, Environmental Encoding, Neural Oscillations, Beamforming

## Abstract

Our ability to navigate in a new environment depends on learning new locations. Mental representations of locations are quickly accessible during navigation and allow us to know where we are regardless of our current viewpoint. Recent fMRI research using pattern classification has shown that these location-based representations emerge in the retrosplenial cortex and parahippocampal gyrus, regions theorised to be critically involved in spatial navigation. However, little is currently known about the oscillatory dynamics that support the formation of location-based representations. We used MEG to investigate region-specific oscillatory activity in a task where participants could form location-based representations. Participants viewed videos showing that two perceptually distinct scenes (180° apart) belonged to the same location. This “overlap” video allowed participants to bind the two distinct scenes together into a more coherent location-based representation. Participants also viewed control “non-overlap” videos where two distinct scenes from two different locations were shown, where no location-based representation could be formed. In a post-video behavioural task, participants successfully matched the two viewpoints shown in the overlap videos, but not the non-overlap videos, indicating they successfully learned the locations in the overlap condition. Comparing oscillatory activity between the overlap and non-overlap videos, we found greater theta and alpha/beta power during the overlap relative to non-overlap videos, specifically at time-points when we expected scene integration to occur. These oscillations localised to regions in the medial parietal cortex (precuneus and retrosplenial cortex) and the medial temporal lobe, including the hippocampus. Therefore, we find that theta and alpha/beta oscillations in the hippocampus and medial parietal cortex are likely involved in the formation of location-based representations.

---

Our ability to navigate in a new environment relies on learning its spatial layout by forming location-based representations. Location-based representations allow us to locate ourselves in space irrespective of the current heading direction and are thought to be formed by binding distinct viewpoints of a single location into a more coherent viewpoint-independent representation. Recent research using pattern classification on fMRI data has found that specific regions within a spatial navigation network support location-based representations (Baumann & Mattingley, 2021; Marchette et al., 2014; Robertson et al., 2016; Vass & Epstein, 2013). In particular, these representations can emerge in medial temporal and medial parietal regions following a single learning session (Berens et al., 2021). Despite evidence for the emergence of location-based representations, we know less about their formation. Specifically, we do not know the oscillatory dynamics that support their formation. Despite the known role of theta and alpha/beta oscillations in memory and spatial navigation (Buzsáki & Moser, 2013; Hanslmayr et al., 2016; Nyhus & Curran, 2010; Parish et al., 2018), they have yet to be specifically related to the formation of location-based representations in humans. As such, we assessed the role of region-specific oscillatory dynamics during at task that allowed for the formation of location-based representations.

Spatial navigation is supported by a network of brain regions, including the medial temporal lobes (and hippocampus) and medial parietal regions (Baumann & Mattingley, 2021; Bicanski & Burgess, 2018; Epstein, 2008; Epstein & Baker, 2019). In the medial temporal lobes, a wide range of neurons have been found in rodents that respond to specific features of the spatial environment during navigation. The archetypal spatial neuron in the hippocampus is the *place cell*, a neuron that fires when the animal is in a specific location, irrespective of the heading direction of the animal (O’Keefe & Dostrovsky, 1971). Thus, place cells are an example of a location-based viewpoint-independent representation in the hippocampus. Other neurons that respond to the animal’s heading direction (Taube et al., 1990), proximity to spatial boundaries (boundary vector cells; (Lever et al., 2009; O’Keefe & Burgess, 1996; Solstad et al., 2008), and to multiple locations in a regular hexagonal pattern (grid cells; Hafting et al., 2005), as well as other spatially-tuned neurons, have also been shown (see e.g., Hartley et al., 2014 for a review). This range of spatially tuned neurons in the medial temporal lobes is thought to support a ‘cognitive map’ of the environment, allowing the organism to effectively navigate in a flexible manner (O’Keefe & Nadel, 1978). Computational models propose that the allocentric representations in the medial temporal lobe are translated into more egocentric navigationally-relevant representations in the parietal lobes, and precuneus in particular, with the retrosplenial cortex acting as a key region in this translation process (Alexander et al., 2023; Bicanski & Burgess, 2018). Thus, the learning of location-based representations is likely to involve both parietal and medial temporal regions, translating more egocentric ‘viewpoints’ into more coherent location-based representations.

Location-based representations have also been observed in the wider spatial navigation network. Using fMRI representational similarity analysis (RSA), representations for known real-world locations have been shown in the parahippocampal gyrus and retrosplenial cortex (Marchette et al., 2014; Steel et al., 2023; Vass & Epstein, 2013). Two recent studies used a novel paradigm to study the formation of these location-based representations using panoramic 2D images of real-world locations (Berens et al., 2021; Robertson et al., 2016). The panorama was cropped to derive scenes that were 180° apart, resulting in two viewpoints of the same location that were perceptually distinct, such that a naïve viewer would not be able to determine if they were from the same location or from two different locations. Participants watched videos panning across the whole panorama, providing an opportunity for the participants to sample shared information between the viewpoints (for overlap videos) and integrate them to form a coherent location-based representation of that panorama. In Berens et al. (2021) participants also watched videos that panned between two viewpoints from different locations (non-overlap videos), where they were not able to form a location-based representation. Participants were able to behaviourally match endpoints in the overlap condition, importantly only after watching the videos, but they were unable to match endpoints in the non-overlap condition. Using this manipulation, they found neural evidence of location-learning in the parahippocampal and retrosplenial cortex. Multivariate patterns of BOLD activity related to viewing two viewpoints from the same location were more similar after, relative to before, watching the overlap videos (no increase in similarity was seen for viewpoints presented in non-overlap videos).

Berens et al. (2021) also assessed univariate BOLD differences between the overlap and non-overlap conditions during the video watching period. This revealed a significantly greater BOLD response to the overlap relative to the non-overlap condition in medial prefrontal regions, and reduced activity in the overlap relative to the non-overlap condition in parahippocampal and retrosplenial cortex (https://neurovault.org/images/115016/). These results are somewhat difficult to interpret, and do not provide satisfactory insight into the neural processes involved in the formation of location-based representations. In particular, given the key role that oscillatory dynamics play in both memory and spatial navigation (Buzsáki, 2002; Buzsáki & Moser, 2013), they do not provide any information about the region-specific oscillations that support the formation of location-based representations.

Both theta (4-8Hz) and alpha/beta (8-20Hz) oscillations have been broadly implicated in spatial navigation and memory (Buzsáki & Moser, 2013). Theta oscillations are theorised to facilitate long-range communication between the hippocampus and neocortex via the striatum and septal nuclei (Arnulfo et al., 2015; Bzymek & Kloosterman, 2023; Gloveli et al., 2005; Hangya et al., 2009; Nyhus & Curran, 2010) in the service of navigation and memory. Neurons in the rodent hippocampus fire in theta frequencies during translational movement (Vanderwolf, 1969) and hippocampal place cells fire at different phases of the theta oscillation, known as theta phase precession (Skaggs et al., 1996). Hippocampal theta phase is also considered important for encoding and retrieval states of episodic memory (Hasselmo et al., 2002). Disrupting the theta rhythm in the rodent hippocampus was found to cause poor performance in spatial memory tasks, while reviving hippocampal theta improved performance (McNaughton et al., 2006). Therefore, there is causal evidence that rodent hippocampal theta oscillations are critical for spatial navigation and memory.

In humans, both invasive (iEEG/ECoG) and non-invasive (EEG/MEG) measurements of electrophysiological activity have implicated the theta rhythm in episodic memory and spatial navigation (Herweg et al., 2020; Nyhus & Curran, 2010). Specifically, changes in theta oscillatory power have been observed on the scalp during learning of specific items (words/pictures/scenes; Fellner et al., 2013, 2019; Guderian et al., 2009; Hanslmayr et al., 2009, 2011; Khader et al., 2010; Osipova et al., 2006), lists of items (Fellner et al., 2016; Griffiths et al., 2016; Meeuwissen et al., 2011; Sederberg et al., 2006), and during associative learning (Backus et al., 2016; Caplan & Glaholt, 2007; Crespo-García et al., 2016; Griffiths et al., 2016; Joensen et al., 2023; Staudigl & Hanslmayr, 2013; Summerfield & Mangels, 2005). Theta power changes have also been found when participants learn spatial layouts as they combine sensory input in new environments compared to when they freely explore a new spatial layout in the absence of an overt task (Chrastil et al., 2022; Du et al., 2023; Kaplan et al., 2012; Pu et al., 2017). Therefore, a large body of evidence indicates that theta power changes observed on the scalp are important for both non-spatial (episodic) and spatial memory formation.

Theta power changes during memory encoding often localise to the medial temporal lobes (among other structures), including the hippocampus and parahippocampal regions (Backus et al., 2016; Crespo-García et al., 2016; Fellner et al., 2016, 2019; Griffiths et al., 2016; Hanslmayr et al., 2011; Staudigl & Hanslmayr, 2013). Note, this theta effect is likely distinct from frontal mid-line theta, found during cognitive control and working memory tasks, which typically localises to medial prefrontal regions (Chrastil et al., 2022; Du et al., 2023; Summerfield & Mangels, 2005; Zuure et al., 2020). Thus, a large body of evidence exists for the role of theta in forming associations between information in both spatial and non-spatial contexts. We therefore predicted that theta rhythms would be implicated in the formation of location-based representations that involves associating different viewpoints from the same location.

Oscillations at alpha and low beta frequencies (∼8-20Hz) are also implicated in memory formation and are found in posterior sites that localise to parieto-occipital regions when scenes/objects are encoded (Jensen & Mazaheri, 2010). Changes in alpha/beta oscillatory power are often thought to reflect functional inhibition, where increases in alpha power are observed in task-irrelevant brain regions, and are thought to reflect inhibition of these regions, and decreases in alpha power are found in task-relevant regions, and are thought to reflect release from inhibition. The oscillatory sync/de-sync model of memory formation (Hanslmayr et al., 2016; Parish et al., 2018) argues that stimulus-specific information is represented by reductions in alpha/beta power in the neocortex, such as visual information represented in parieto-occipital regions (Griffiths et al., 2019; Jensen & Mazaheri, 2010; Klimesch, 2012). This stimulus information then travels to the hippocampus, where hippocampal theta power increases allow for mnemonic information to be integrated into a coherent memory trace via long-term potentiation. More recently, alpha oscillations have also been associated with micro-saccades and gaze towards to-be-encoded stimuli indicating that changes in alpha/beta power, especially in the visual cortex, could also be involved in memory formation (Popov & Staudigl, 2023; Staudigl et al., 2017). Therefore, there are theoretical and computational models backed by empirical evidence that indicate changes in theta and alpha/beta oscillations (∼4-30Hz) are involved in associative memory formation.

Despite the wealth of data implicating theta and alpha/beta in spatial navigation and memory, no study to date has focussed on these oscillations specifically during the formation of location-based representations. Further, no study has attempted to source-localise the oscillations supporting the formation of location-based representations. We therefore used MEG to measure electrophysiological activity related to the formation of location-based representations because it is temporally precise and provides a direct measure of changes in neural oscillations. Further, we aimed to estimate neural activity arising from the hippocampus and medial parietal structures, which are traditionally considered to be challenging to record from because of the depth of these regions relative to the scalp, and because the complex gyri folds in these regions could cancel out any signals before they emerge at the scalp (Pu et al., 2018). Although these problems do exist, mounting evidence indicates that hippocampal activity can be localised using advanced beamforming algorithms, especially with MEG data (Guitart-Masip et al., 2013; Korczyn et al., 2013; Pizzo et al., 2019; Pu et al., 2018; Ruzich et al., 2019). Therefore, using beamformers on MEG data, we predicted changes in theta (4-8Hz) (and potentially higher frequency alpha/beta (8-30Hz)) oscillatory activity from the hippocampus and medial parietal structures during the formation of location-based representations.

To test this, we adapted Berens et al.’s (2021) paradigm for MEG. Participants watched the overlap and non-overlap videos while we recorded neural activity. We tracked the formation of location-based representations behaviourally by assessing their ability to match viewpoints from the same location before and after watching the videos. We use the term “location-based representation” in this task as a short-hand for a mnemonic representation that allows participants to infer which viewpoints are from the same location after watching the panoramic videos, without making explicit claims about the precise nature of the representation (i.e., whether they relate to true allocentric representations of location such as place cells). We predicted that neural oscillatory power in the theta band would be greater when participants watched overlap videos, when location-based representations can be formed (Berens et al., 2021; Robertson et al., 2016), compared to non-overlap videos, when location-based representations cannot be formed. Given evidence for alpha/beta oscillations in memory (Hanslmayr et al., 2016; Parish et al., 2018), we also assessed power in frequency bands other than theta in a more exploratory manner. We also attempted to find evidence for the successful formation of location-based representations by comparing neural activity at encoding (watching overlap videos) based on whether they were subsequently remembered vs. forgotten (subsequent memory effect). This subsequent memory effect is often characterised by changes in hippocampal theta oscillations (Hanslmayr et al., 2016; Joensen et al., 2023; Parish et al., 2018; Voss et al., 2017), although the specific direction (increase or decrease) of change is dependent on several factors (Herweg et al., 2020). Thus, we predicted hippocampal theta power changes while participants watched overlap videos that were subsequently remembered compared to forgotten.

## Methods

### Participants

Thirty-eight University of York students aged 18 to 34 years (*M =* 23.33 years, *SD* = 4.24, 32 females) participated in this study in exchange for course credits or £20. All participants indicated that they were right-handed, had normal or corrected-to-normal vision, had no prior or existing neurological or psychological illnesses, and were not on any psycho-active drugs. A total of eight participants were excluded due to issues during MEG acquisition (3), and extremely noisy MEG recordings (5). The final sample size was 30. Ethics approval was granted by the York Neuroimaging Centre’s (YNiC) ethics committee. Informed consent was obtained from all individual participants included in the study. Participants consented for anonymised data to be available on online repositories and to be published in peer-reviewed journals.

### Stimuli and Design

Our experimental paradigm was based on Berens et al., (2021). The stimuli were created from twelve panoramic images of locations in different UK cities. Each panorama spanned a 210° horizontal field-of-view. The two endpoints of each panorama were extracted and displayed 30° of extreme ends of the panoramas. As such, naïve participants would not be able to accurately match two endpoints from the same location (see Berens et al., 2021). These endpoints were used in the endpoint-matching tasks conducted both before and after a learning task. See Figure 1.A-B for an illustration of the stimuli used in the study.

**Figure 1.**
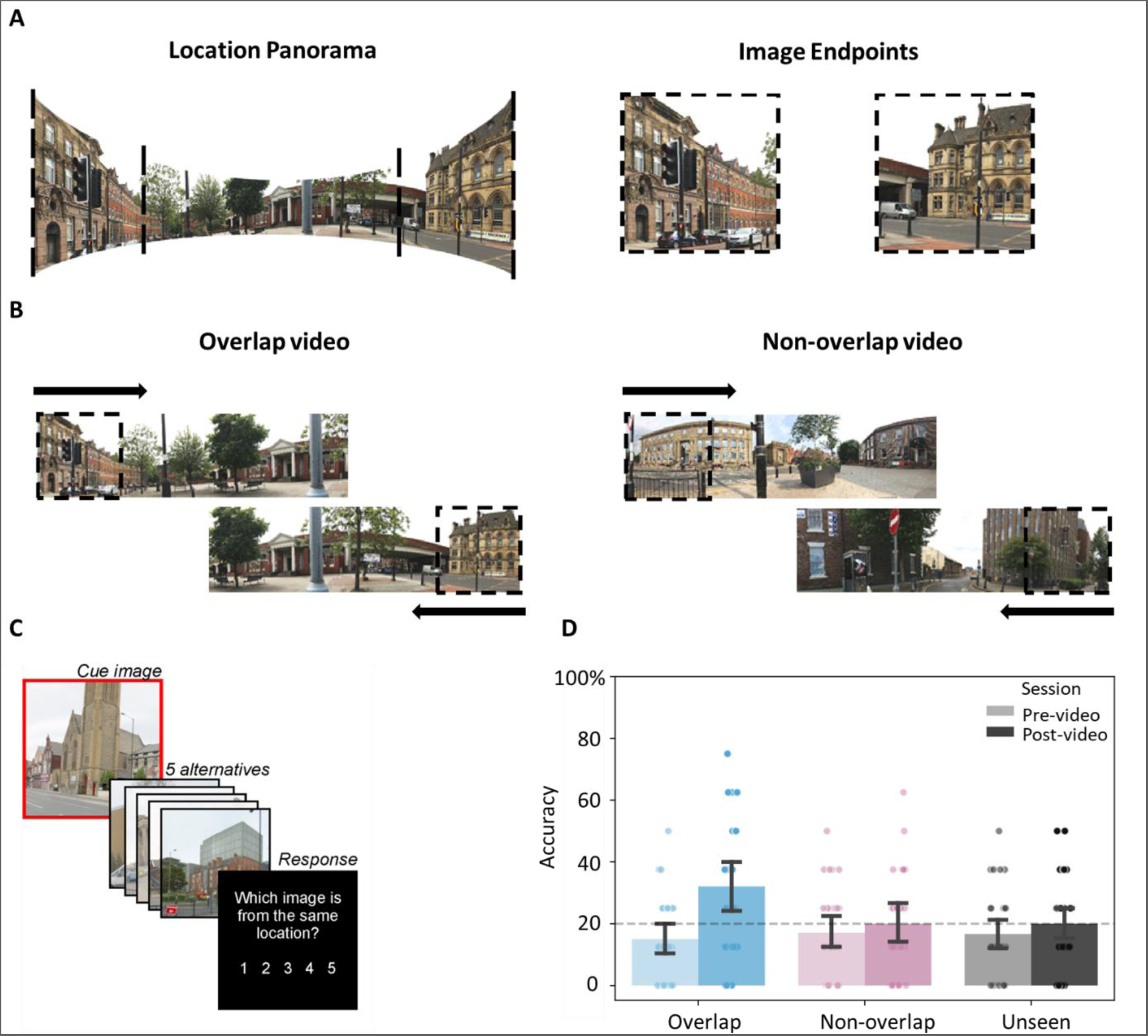
Stimuli used in the experiment and results from the behavioural tasks. Panels A-C are adapted from Berens et al., (2021). (A) An example panorama of a location with the two endpoint images marked with dotted lines. The whole panorama was never shown to the participants, like previous experiments (Berens et al., 2021; Robertson et al., 2016). (B) Videos participants watched in the MEG scanner. Both overlap and non-overlap videos panned from one endpoint to the centre, then switched to the other endpoint and panned to the centre. The central region is from the same panorama in overlap videos, but from different panoramas in non-overlap videos. Overlap videos therefore allowed for the formation of location-based representations. (C) Example trial in the endpoint-matching task. The target image was bordered in red and followed by presentation of 5 lures. Participants chose which lure belonged to the same location as the target. (D) Endpoint-matching accuracy compared between pre- and post-video endpoint-matching tasks grouped by video condition. Scatter dots reflect data from individual participants. Error bars represent 95% confidence intervals. Chance level for 5-AFC is 20%, represented by a dashed line.

In the learning task, participants viewed videos panning from one endpoint to the midpoint of a panorama, then switching to another endpoint and panning to the midpoint. In the overlap condition, participants viewed camera pans of the same panoramic scene, whereas in the non-overlap condition, participants viewed camera pans from two different panoramas. For instance, in the overlap condition, the left-to-centre sweep would be from the left endpoint of a location panorama and the right-to-centre sweep would be from the right endpoint from the same panorama. Therefore, there was clear visual overlap between both sweeps and participants could infer that the two endpoints belonged to the same location. However, in the non-overlap condition, the left-to-centre sweep would be from the left endpoint from one location and the right-to-centre sweep would be from a different location. Therefore, there was no visual overlap in non-overlap videos and participants could not infer which location the two endpoints belonged to - they could only infer that the two endpoints did not belong to the same location. The overlap condition was our key manipulation, where participants could form more coherent location-based representations. Non-overlap videos were the control condition, where participants could form associations between the two endpoints but critically were not able to form a more coherent location-based representation.

Eight panoramic scenes (from a total of 12) were used in the videos: Four in the overlap and four in the non-overlap condition. Endpoints from the other four panoramas were unseen in the video task but were used as behavioural controls in the pre- and post-learning tasks. The assignment of endpoints to each condition was fully counterbalanced. The starting location of the video pan (left or right endpoint of the panorama) was counterbalanced both across and within participants using the following rules: The 12 panoramic images were divided into 4 factions (sets) that had 3 panoramas (A, B, C) and 6 sets of endpoints (A1, A2, B1, B2, C1, C2) in each faction. The 6 endpoints were matched in the video-watching task such that A1-A2 were in the overlap videos, and belonged to the same panorama, B1-C2 were in the non-overlap videos, and C1 and B2 were unseen in the video-watching task (for one of the counterbalancing sets).

Both end-to-centre pans were repeated twice in each video to ensure that the visual overlap was easily detectable, in line with Berens et al., (2021). Each end-to-centre pan lasted for 5s: The endpoint was shown for 1s, the pan from the endpoint to the centre took 3s, and the midpoint was shown for 1s. Each video had four pans. The first pan was from left-to-centre, the second was from right-to-centre, third was again from left-to-centre of the same panorama, and the final pan from right-to-centre of the same panorama. Therefore, there were 4 total pans lasting 5s each equating to 20s for each video. For counterbalancing purposes, half of the videos started with a left-to-centre pan and the other half of the videos started with a right-to-centre pan (see Figure 2 for an illustration).

**Figure 2.**
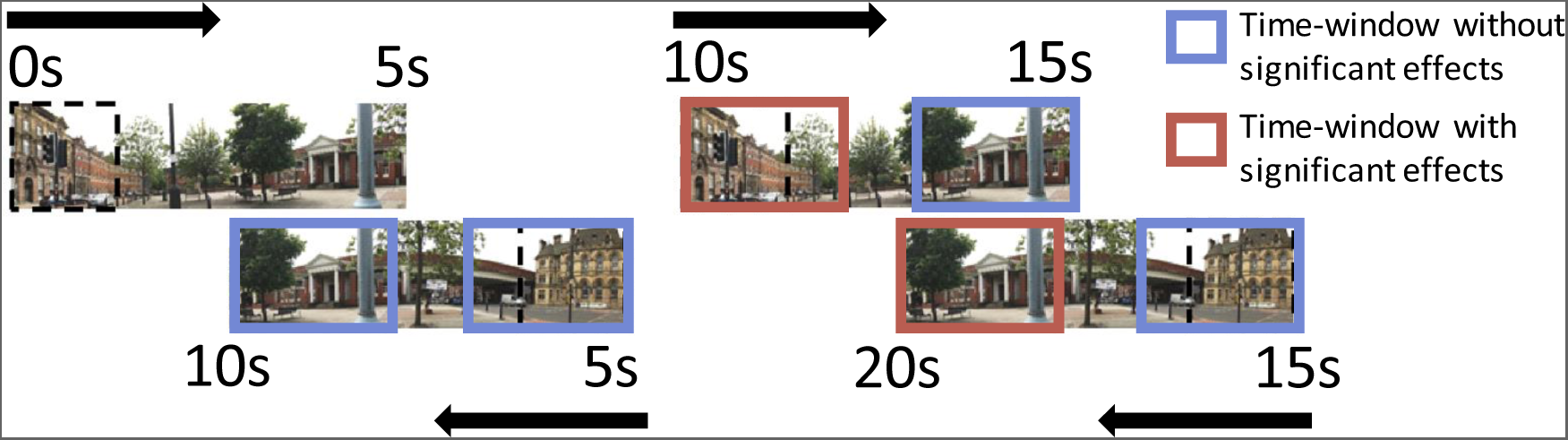
Illustration of one trial in the video-watching learning task. Each trial consisted of two presentations of the location panorama, and 2 camera pans for each presentation, leading to a total of 4 camera pans. Each camera pan lasted for 5s, leading to 20s in each trial. For half of the trials, camera pans were in this order: left-to-centre, right-to-centre, left-to-centre, right-to-centre. The other half of the trials started with right-to-centre camera pans and alternated with left-to-centre pans. The video jumps to the other camera pan after completing the previous camera pan. For the MEG oscillation analysis, we chose three time-windows immediately after a camera pan began, (5 to 6.5s, 10 to 11.5s, and 15 to 16.5s), at end-point onset; and three time-windows around the mid-points of the camera pans (8 to 9.5s, 13 to 14.5s, and 18 to 19.5s), starting at the time-points where the shared location information starts becoming apparent. The solid borders depict these chosen time-windows. We found significant changes in neural activity during overlap vs. non-overlap video-watching in the 10-11.5s and 18-19.5s time windows, and these are depicted by pink borders. We did not find significant effects in the other time-windows, which are depicted by blue borders. The black dashed border indicates an endpoint image. Non-overlapping videos also followed the same exact presentation order.

### Procedure

We adapted Berens et al.’s (2021) experimental paradigm for MEG data acquisition. Similar to Berens et al. (2021), participants first completed a behavioural task that assessed their ability to match endpoints from the same location (outside the MEG scanner). Then, participants entered the MEG scanner, where they were presented with each endpoint before and after the learning phase (oddball detection task). In the learning phase they watched the overlap and non-overlap videos (video-watching learning task). Outside the MEG scanner, participants then completed the same endpoint matching task they completed before entering the scanner, to assess learning.

#### Behavioural pre- and post-scanner tasks

This task was conducted both pre- and post-video-watching to assess if participants were able to accurately match two endpoints from the same location after (but not before) watching the panoramic videos. On each trial, participants first viewed one endpoint for 3s that was surrounded by a red box. Then, five other endpoints were sequentially presented with a number alongside (i.e., 1-5; 2s per endpoint; 500ms ISI). Out of the five images, one image was from the same location as the cue (red bordered endpoint), and the other four endpoints were from different locations, but from the same counterbalancing set (faction). The presentation order of the five endpoints was random. Participants had to indicate the endpoint that belonged to the same location as the cue (red bordered endpoint) via button press. See Figure 1.C for an illustration of this task.

A secondary endpoint-matching task was conducted after the post-scanning task described above. The trial structure was identical, but the instructions were different. Participants were asked to match endpoints that were shown in the same video, rather than match endpoints from the same location (as they did in the previous task). Unseen endpoints were not presented in this task as targets but were presented as foils. Only endpoints that belonged to overlap and non-overlap videos were presented as cues in this task. Note that this is different because participants could still match endpoints from non-overlap videos in this task as they appeared in the same video but would still not know which endpoints belong to those respective locations.

#### Oddball detection task

Participants completed an oddball detection task where they viewed presentations of endpoints in the scanner both before and after the video-watching learning task. On one trial, a fixation cross appeared for a jittered duration centred on 2s (2.25s to 1.75s), followed by a scene that lasted for 3s. Eight blocks of 24 endpoints were presented and the order of endpoint-presentation was pseudo-randomised both within and across blocks. Therefore, each endpoint was presented 8 times. Three trials were randomly chosen in each block to be oddball trials. Participants pressed a button to indicate that they saw an oddball image. The oddball stimulus was an endpoint with three small red dots (3-pixel radius) superimposed in random locations on the image.

This task was used to run multivariate pattern analyses to identify representations of individual endpoints. However, we were not able to identify patterns for individual scenes reliably, and as such were not able to perform analyses to assess changes in patterns as a function of learning (as in Berens et al., 2021). We report these analyses in the Supporting Information for completeness.

#### Video-watching learning task

The video-watching task was our main task of interest, as we were interested in the oscillations that underpinned learning of location-based representations. See Figure 2 for an illustration of one trial in the video-watching task. We collected MEG recordings as participants viewed the overlap and non-overlap videos. On one trial, a fixation cross first appeared for 2s (jittered between 1.75s to 2.25s), followed by a video that lasted 20s. After a 2s interval, participants were asked to indicate whether the videos were from the same location or from different locations to ensure that they were paying attention to the visual overlap across segments. Participants used the left/right keys on a response box to make these decisions and had 3s to respond. Then, a 2s inter-trial-interval was followed by the next trial. Importantly, participants were asked to only respond when the text appeared on the screen asking for a response, to avoid motor-related neural activity during video-watching. Each video was repeated three times across the task. Each video was presented a single time within 3 blocks, with presentation order randomised in each block.

### MEG data acquisition and processing

We measured neural activity with a 4D Neuroimaging Magnes 3600 whole-head 248-sensor magnetometer-based MEG. Four sensors were permanently offline, so we acquired data from 244 sensors for all participants at a 1000Hz sampling rate. Participants’ head-shape for source-localisation was attained using a Polhemus Fastrak Digitisation System. MEG data were pre-processed and analysed using a combination of the FieldTrip toolbox (Oostenveld et al., 2011) and self-written MATLAB code.

Raw MEG recordings were first band-pass filtered between 0.1Hz and 70Hz using two-pass butterworth filters. Notch filters were applied at 50Hz (UK mains frequency) and 60Hz (projector refresh rate). Further details of analyses and results of the oddball detection task are detailed in the Supporting Information. We focus on the video-watching task as we were primarily interested in neural oscillations during learning. Further, no reliable multivariate patterns were found in the oddball detection task.

The filtered continuous data from the video-watching task were segmented into epochs of 24s, from - 2s pre- to +20s post-stimulus onset. Trial-by-trial information about stimulus type and participant responses was then added to the FieldTrip data structure. Noisy epochs were rejected by visual inspection: As a first pass, experimenters looked for any specifically noisy trials including scanner drifts/jumps and extreme movements. Remaining epochs were concatenated and submitted to an Independent Component Analysis (ICA) using *runica* in FieldTrip with default parameters. Note that ICA was conducted by band-pass filtering the dataset between 1 to 40Hz and resampling to 200Hz, which is argued to improve ICA performance (Ferrante et al., 2022). Importantly, artefact-rejection occurred on the original data set, which was filtered between 0.1 to 70Hz. Components reflecting eye-movements and heartbeats were identified by visual inspection of component scalp topographies and time-courses. These components were discarded by back-projecting all but these components to the data space. Corrected data were low-pass filtered to 40Hz (two-way butterworth). Finally, any remaining noisy epochs were removed based on visual inspection and were baseline-corrected against the −200 to 0ms prestimulus period. The data were downsampled to 250Hz before further analysis.

After pre-processing, an average of 11.30 trials (*SD* = 1.24, range 7 to 12) in the overlap condition and 11.06 trials in the non-overlap condition (*SD* = 1.17, range 8 to 12) remained and were used for further analyses.

#### Source reconstruction of MEG activity

Observed sensor-level data was reconstructed in source-space using Linear Constrained Minimum Variance (LCMV) beamformers. A standard MNI structural T1 image from the FieldTrip toolbox was aligned to each individual’s headshape obtained immediately before participants entered the MEG scanner. The template structural scan was first co-registered to the fiducials and transformed to fit the headshape automatically. The experimenter then visually verified fit and manually resized each co-registered image to ensure good fit between the scan and the digitised headshape. The co-registered image was resliced to align voxel positions in 3D space to the individual’s headshape and segmented to isolate brain tissue from skull and skin. The segmented image was used to create positions of 3294 grid points spaced 10mm inside the brain. The lead-field matrix (forward model) was then computed, which included information about the forward projections from each grid point to scalp sensors. The sensor level data, source model, and forward model were input into the beamforming algorithm, with fixed dipole orientations and 5% lambda regularisation. The beamformer output resulted in time-series signals from −2s to +20s centred around video onset in each of the 3294 voxels. These signals were then decomposed into time and frequency dimensions to derive oscillatory power using aforementioned parameters.

#### Time-frequency decomposition

Single-trial epochs were convolved with complex Morlet wavelets to derive oscillatory power of the MEG signal across time and frequency, using MATLAB code adapted from Denis et al. (2021). Twenty-seven wavelets with centre frequencies ranging between 4-30Hz were used. Both the wavelet width and frequency steps were logarithmically spaced in order to distribute power equally across all theoretically important frequency bands (Theta: 4-8Hz, Alpha: 8-12Hz, and Beta: 8-30Hz). Linear spacing of frequencies would have prioritised higher frequency bands (Beta; 13 steps) rather than lower frequency theta (4 steps) and alpha (4 steps) bands. To remove edge artefacts, epochs were truncated to −1.75 to 19.5s. A pre-stimulus baseline period of −1 to −0.5s was used to normalise the oscillatory power to decibels (dB), because using baselines close to stimulus-onset could lead to bleeding of stimulus-related activity into the baseline period. The decomposition parameters were identical for both sensor-space and source-space data.

### MEG data analysis

Our primary analysis was a pairwise comparison of oscillatory activity between overlap and non-overlap videos. Non-parametric cluster-based permutation tests (N_perms_ = 1000, two-tail correction = ‘prob’ which multiples the uncorrected probability by a factor of two before setting the α = .05 threshold) as implemented in FieldTrip were used to detect statistically significant differences between these two conditions in both sensor and source-space, across time and frequency.

#### Sensor-level analysis

The cluster tests were first conducted in sensor-space (with a minimum of 3 neighbouring sensors required to form a cluster). We could not test for effects across all 20s of video watching in sensor-space because of RAM limitations (but this is addressed later). Therefore, we divided the 20s epoch into 1.5s long segments (the maximum time-window we could test with 1000 permutations) of interest around endpoint and midpoint onsets. We expected that participants would most likely be able to distinguish the endpoints were from same or different locations at these points in the videos. Hence, we expected oscillatory power involved in learning to be maximal between the overlap and non-overlap videos during the end- and mid-points of the videos.

We specified 3 time-segments starting at endpoint-onset (5 to 6.5s, 10 to 11.5s and 15 to 16.5s), and three segments around mid-point onset (8 to 9.5s, 13 to 14.5s, and 18 to 19.5s), see Figure 2 for an illustration of where these time-segments lie in the video. We specified these time-windows because they either represented the to-be-bound information (the endpoints) or depicted shared spatial information (mid-points) allowing binding between endpoints into a location-based representation. So, we expected the binding process that forms location-based representations to occur during these time-windows, which would be reflected by changes in neural oscillatory activity that we intended to find. The final 500ms of mid-point presentation could not be analysed in any situation. In the first two mid-point segments, evoked activity from the next endpoint-onset bled into preceding timepoints of mid-point-offset. The final 500ms included edge-artefacts arising from time-frequency decomposition.

Finally, we attempted to detect clusters across the whole 20s of video watching that we may have missed. We chose sensors that had the maximal t-value within significant clusters from the focal time-window analysis. Then, we ran cluster-based tests to detect significant clusters from 0 to 19.5s and 4 to 30Hz in each sensor. Note, this analysis can only be used to reveal significant effects at timepoints outside of our initial time-windows of interest, given the sensors were selected based on analyses from these time-windows.

#### Whole brain analyses

We then assessed regions in the brain that contributed to any statistically significant sensor-space differences. Oscillatory power in each voxel in the brain was averaged over the time-frequency regions spanned by significant clusters found in sensor-space. We computed an averaged oscillatory power value for each voxel across the whole brain and used cluster-permutation pairwise tests to find differences in averaged oscillatory power between overlap and non-overlap videos across all voxels. The Brainnetome Atlas (Fan et al., 2016) was used to identify regions with statistically significant whole-brain clusters. This analysis therefore revealed the brain regions most likely to be producing the sensor-level power differences.

#### ROI analysis

We ran targeted ROI analyses with two goals in mind. First, we wanted to reliably test for any clusters at time-points other than the segments chosen for sensor-space analyses. Second, although the whole-brain analyses revealed the brain regions involved in sensor-level effects, the distribution of these effects across time and frequency within specific brain regions was relatively unknown. Therefore, we tested for the extent of differences in oscillatory power across time-frequency ranges in each ROI and time-segments where we found sensor-level effects.

We chose regions of interest that were theorised to be critical for supporting location-based representations (see Alexander et al., 2023; Bicanski & Burgess, 2018; Byrne et al., 2007): the precuneus, retrosplenial cortex, and right and left hippocampi. The hippocampus is also critical for integrating episodic and spatial information (e.g., Backus et al., 2016; Buzsáki & Moser, 2013; Horner et al., 2015).

LCMV beamformers are more likely to generate illusory effects in either the right or left hippocampus as the algorithm fails to separate correlated sources (O’Neill et al., 2021). Therefore, any differences in power between conditions would likely be (falsely) lateralised in either the left or right hippocampus, and averaging over the two hippocampi could reduce our power to detect any true effects. Therefore, we ran our tests separately in the right and left hippocampus. Crucially, we did not make inferences about the lateralisation of any effects we found but were simply attempting to increase statistical power by running our analysis separately in the right/left hippocampus. Note that the precuneus and retrosplenial cortex were combined across left and right hemispheres as they were physically proximal and do not suffer from the illusory lateralisation found in the physically distant but functionally connected hippocampi.

Voxels belonging to each ROI were identified by interpolating grid-points with the Brainnetome atlas (Fan et al., 2016) using FieldTrip functions. We averaged oscillatory power across all voxels from each ROI for each data-point from 0 to 19.5s and 4 to 30Hz. Cluster-based permutation tests were conducted to detect differences in oscillatory power across time and frequency in each ROI across (i) 0 to 19.5s and (ii) in time-segments of interest from the sensor-level analysis where we found significant clusters (10 to 11.5s and 18 to 19.5s, see Figure 2).

#### Subsequent memory effect

In a complementary analysis, we tested for subsequent memory effects in each of the four ROIs. We split overlap videos based on whether participants were accurate or inaccurate in the subsequent behavioural scene-matching test. We then extracted average power from the subsequently accurate and inaccurate videos by defining time-frequency boundaries found in the overlap vs. non-overlap comparison in each ROI. Locations where participants could match only in one direction (endpoint B cued by endpoint A, or vice-versa) were considered to be subsequently remembered in order to increase trial numbers and statistical power (given the above chance but relatively poor performance post-learning in the overlap condition). A frequentist pairwise t-test was conducted between subsequently accurate and inaccurate videos from the video-watching task in each ROI that had significant overlap vs. non-overlap clusters. We had to exclude data from 6 participants from this analysis as they either remembered or forgot all the locations, and therefore did not have any trials in either one of the conditions. For the remaining 24 participants, 5.63 subsequently accurate (remembered) trials (SD = 2.32, range = 1 to 9) and 5.50 subsequently inaccurate (forgotten) trials (SD = 2.40, range = 1 to 9) remained on average, and were used for further analyses.

## Results

### Behavioural results

We tested our behavioural hypothesis that endpoint-matching accuracy would be greater post- vs. pre-video watching of overlap but not non-overlap videos or unseen endpoints (see Figure 1.D).

Average endpoint-matching accuracy was submitted to a 2 (Session: pre-video, post-video) x 3 (Condition: overlap, non-overlap, unseen) within-subjects ANOVA. The main effect of Condition was not significant (*F*(1,29) = 1.78), but there was a significant main effect of Session: *F*(1,29) = 11.69, *p* = .002, *η^2^_p_* = .29), where endpoint-matching accuracy was greater post compared to pre-video watching. Importantly, a significant interaction between Condition and Session (*F*(1,29) = 5.91, *p* = .005, *η^2^_p_* = .17) was found. Follow up paired t-tests revealed that participants were able to match endpoints more successfully post- vs. pre-videos for overlap (*t*(29) = 4.15, *p* < .001, *BF_10_* = 96.43, *d* = .75) but not for non-overlap (*t*(29) = 0.72, *p* = .47, *BF_10_* = 0.25) and unseen endpoints (*t*(29) = 1.22, *p* = .23, *BF_10_* = 0.38). Furthermore, participants performed at chance level (20%) when matching endpoints from non-overlap videos (*t*(29) = 0.14, *p* = .89, *BF_10_ =* 0.196) and unseen endpoints (*t*(29) = 0.16, *p* = .88, *BF_10_ =* 0.197) but they performed significantly better than chance at matching endpoints from the same location post-videos (*t*(29) = 3.09, *p* = .003, *BF_10_ =* 9.08, *d* = .56). Therefore, participants learnt that two endpoints belonged to the same location after watching overlap compared to non-overlap videos and unseen locations, where they did not learn that two endpoints belonged to the same location. We also analysed the behavioural data using the same GLMMs as in Berens et al. (2021) and found a similar pattern of results (see Supporting Information).

Participants were at chance when matching two scenes from the same video for both overlap and non-overlap videos (*M*_overlap_ = 21.7%, *SD* = .17, *BF_10_* = 0.22, *t*(29) = .53, *p* = .59; *M*_non-overlap_ = 21.7%, *SD* = .19, *BF_10_* = 0.22, *t*(29) = .48, *p* = .63), and performance did not differ between both conditions (*BF_10_* = 0.19, *t*(29) = 0.1, *p* = .99). Participants were hence unable to determine which endpoints were shown in the same video for both overlap and non-overlap videos. It is not clear why participants could not match endpoints for the overlap condition, given they were able to in the main task reported above. It is possible that they did not fully understand the task or were not motivated to perform given it occurred at the end of the experiment. Participants were above chance in the overlap condition in this task in Berens et al. (2021), suggesting that participants can do this task if sufficiently motivated.

### MEG results

We next analysed MEG data collected during video-watching to determine the neural oscillations involved in forming location-based representations.

#### Sensor-level results

We first compared oscillatory power between overlap and non-overlap video-watching in sensor-space and estimated sources in the brain underlying the sensor-space effects (see Figure 3).

**Figure 3.**
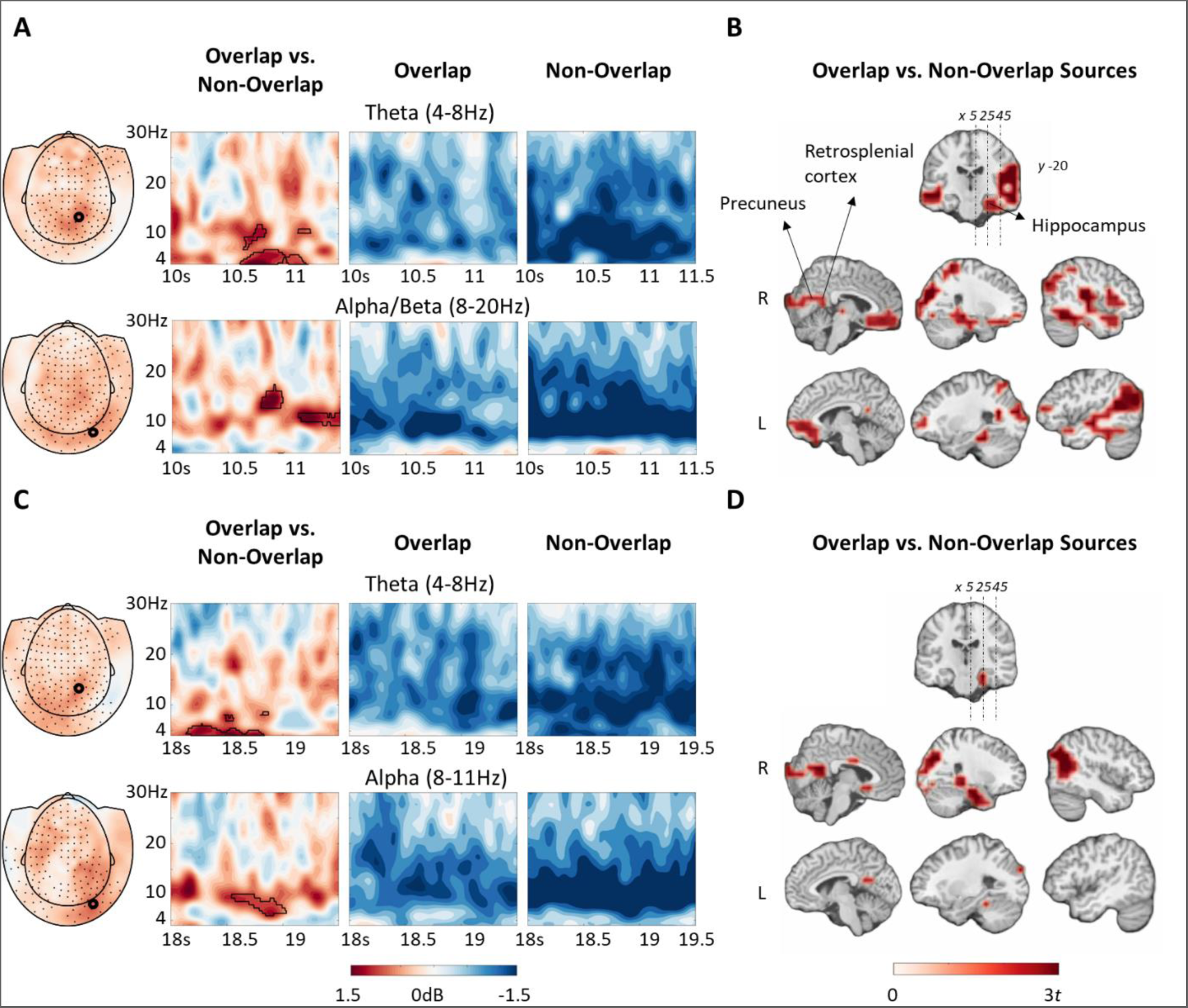
Results from the cluster-based permutation tests in MEG sensor-space and estimated sources of the sensor-space effects. (A) Topographical plots of averaged power across timepoints from the significant cluster in the theta (top) and alpha (bottom) bands. Spectrograms from a posterior sensor (marked on the topographical plot) reflecting baseline-corrected oscillatory power (dB) from 10 to 11.5 seconds post-video onset. Time-frequency data-points that belonged to the cluster are bordered by black lines. (B) Estimated brain sources of the sensor-space effect found during 10 to 11.5s post-video onset between 4 and 20Hz. (C) Topographical plots of averaged power across timepoints from the significant cluster in the theta (top) and alpha (bottom) bands. Spectrograms from a posterior sensor (marked on the topographical plot) reflecting baseline-corrected oscillatory power (dB) from 18 to 19.5 seconds post-video onset. Time-frequency data-points that belonged to the cluster are bordered by black lines. (D) Estimated brain sources of the sensor-space effect found during 18 to 19.5s post-video onset between 4 and 11 Hz.

We found significant clusters in the 10-11.5s and 18-19.5s time-windows. Oscillatory power was greater in theta, alpha, and low beta frequencies from 10.5-11.5s (4-20Hz, *p* = .023) and from 18-19s (4-11Hz, *p* = .03) post-video onset for overlap compared to non-overlap videos. These effects were maximal in posterior sensors but were spread across the whole scalp. No significant clusters were seen during the other time windows of interest (5-6.5s, 8-9.5s, 13-14.5s, 15-16.5s). Note, these effects are cluster corrected within each time-window but are not corrected for multiple comparisons across the 6 time-windows of interest. Neither the 10.5-11.5s or 18-19s effects survive a stringent Bonferroni correction (p<0.05/6 time-windows of interest).

Then, we tested for differences in oscillatory power between overlap and non-overlap videos at time-points other than the a-priori chosen time-windows. To do this, we chose one sensor from the 10-11.5s and 18-19.5s each that had the maximal t-value from the statistically significant cluster showing power changes for the overlap compared to non-overlap condition. Then, we ran a cluster-based permutation test from 0 to 19.5s, which spans all time-points in the video-watching task, across 4 to 30Hz, in each of these sensors. We did not find any clusters that were statistically significant (all *ps* > .34). Therefore, the differences in theta and alpha power seen between 10.5-11.5s and 18-19s in the theta and alpha band appear to be specific to those timepoints.

We next estimated sources in the brain that generated the effects we found on the scalp. To estimate the sources underlying the effect found in the 10-11.5s time window (Fig 3.A), we averaged over 10.5-11.5s and 4-20Hz time-frequency data-points for each voxel in the brain. Similarly, we averaged over 18-19s and 4-11Hz for the second cluster (Fig 3.C) for each voxel in the brain. Then, we ran cluster-based permutation tests to determine the extent of the effect across all voxels in the brain and used the Brainnetome atlas (Fan et al., 2016) to identify regions that belonged to the cluster.

Several brain regions, primarily belonging to the default-mode network and medial temporal lobe, contributed to the greater theta/alpha power for overlap compared to non-overlap videos observed on the scalp (cluster correction: 10.5-11.5s, *p* = .007; 18-19s, *p* = .002). Both effects survive a Bonferroni correction for the two time and frequency windows of interest (p<0.05/2). Oscillatory power changes during 10.5-11.5s of video-watching were localised to bilateral superior, middle, and inferior temporal gyri, superior and inferior parietal cortices, and medial and lateral occipital cortices (Fig 3.B). Notably, both the left and right hippocampus, retrosplenial cortices, preuneus, and vmPFC showed significant power differences between overlap and non-overlap videos. A subset of these same regions were involved in the theta/alpha changes found during 18 to 19s of overlap compared to non-overlap video-watching (Fig 3.B), including the Inferior parietal lobe, hippocampus, retrosplenial cortex, and precuneus. Although the effect appears to be isolated to the right hemisphere, we do not make inferences about the lateralisation because the beamformer algorithm can output illusory lateralised effects (O’Neill et al., 2021).

#### ROI analyses

We then investigated oscillatory power changes specifically in the right and left hippocampus, retrosplenial cortex, and precuneus, as these structures are thought to be important for location-based representations (Bicanski & Burgess, 2018). Moreover, the hippocampus is particularly important for episodic and spatial memory formation (Burgess et al., 2002; Herweg et al., 2020). We determined the distribution of oscillatory power changes for overlap vs. non-overlap videos across time and frequency within our ROIs, within the time-windows of video-watching where we found sensor-level effects: 10 to 11.5s and 18 to 19.5s (see Figure 4).

**Figure 4.**
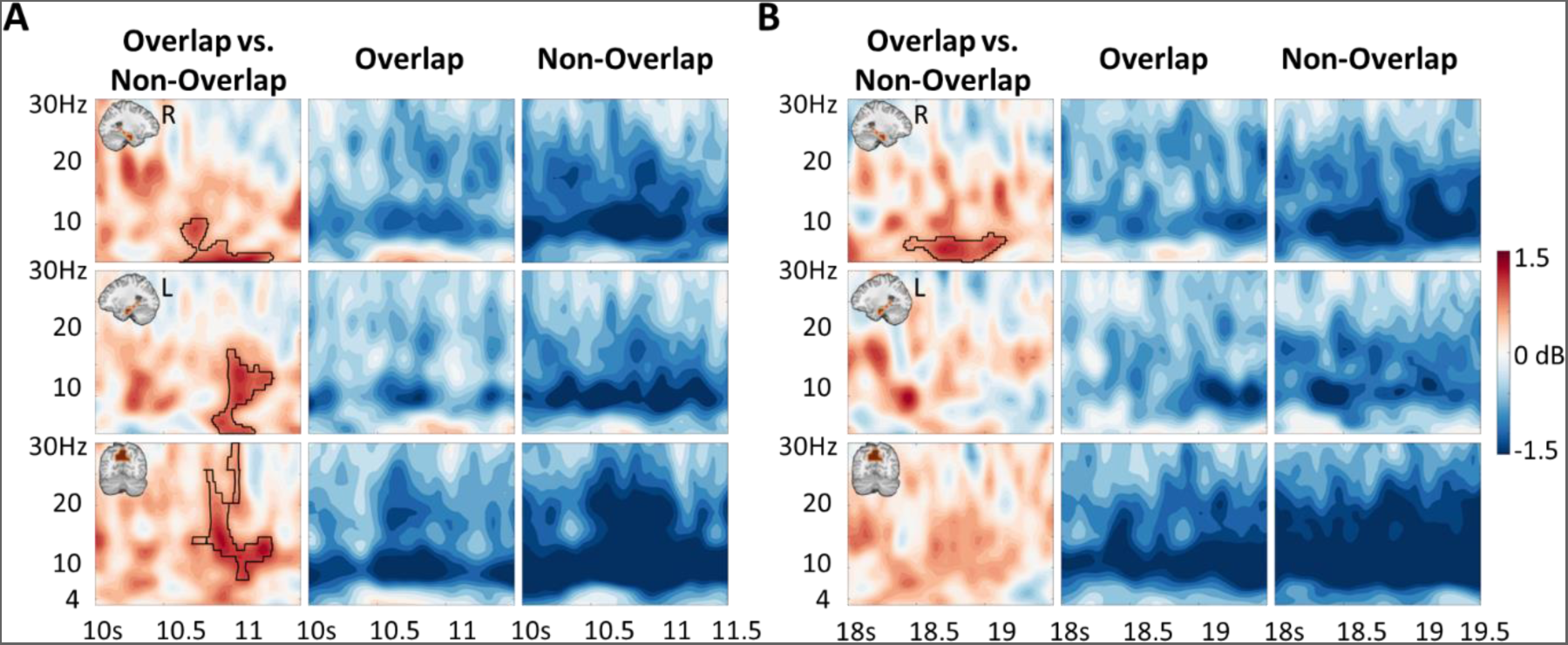
Results from region-of-interest analyses of oscillatory power. (A) Spectrograms depicting oscillatory power (dB) from 10 to 11.5s post-video onset in the right hippocampus (top), left hippocampus (middle), and precuneus (bottom). Time-frequency data-points that belonged to the cluster are bordered by black lines. (B) Spectrograms depicting baseline-corrected oscillatory power (dB) from 18 to 19.5s post-video onset in the right hippocampus (top), left hippocampus (middle), and precuneus (bottom). Time-frequency data-points that belonged to the cluster are bordered by black lines.

In the 10 to 11.5s time window, the power increase for overlap vs. non-overlap videos was in the theta and alpha (4 to 11Hz) bands from 10.5 to 11.5s in the right hippocampus (*p* = .004) and in theta, alpha, and early beta bands (12 to 18Hz) in the left hippocampus (*p* = .02). However, alpha/beta (8 to 28Hz) frequencies dominated in the precuneus (*p* = .045). There were no significant clusters in the retrosplenial cortex during this time window (though we note a cluster in the theta and alpha bands (4 to 11Hz) that was close to the cluster-corrected threshold (*p* = .053). In the 18 to 19.5s time window, in the right hippocampus, the oscillatory power increase for overlap vs. non-overlap videos was in the theta band (4 to 8.6Hz; *p* = .006). There were no significant clusters in the left hippocampus, retrosplenial cortex, or the precuneus in this later time window. Only the right hippocampal effects in the 10.5-11.5s and 18-19.5s time-window survive a stringent Bonferroni correction for multiple comparisons (p<0.05/8, 4 ROIs and 2 time-windows of interest).

We then tested for significant effects across all time-points during video-watching (0 to 19.5s). We focused on the theta (4 to 8Hz) and alpha (8 to 12Hz) bands across 0 to 19.5s (see Figure 5). There were no significant clusters in the alpha band. In the right hippocampus, we found 4 significant clusters indicating greater theta power for overlap vs. non-overlap videos. The two clusters found in previous analyses were also detected in this analysis (10.5 to 11.3s, *p* = .036; 18.5 to 19.1s; *p* = .02). We found two additional clusters from 15.2 to 16.2s (*p* = .047) and 17.11 to 17.55s (*p* = .043). The 15.2 to 16.2s time-window is 200ms post-onset of the second endpoint in time, after the camera pans from the first endpoint at 10s and ends at the central overlapping scene at 14.99s. The 17.11 to 17.55s time-window occurs during the camera pan from second endpoint in time to the final presentation of the central overlapping scene (around 18s). None of these effects survive a stringent Bonferroni correction for multiple comparisons (p<0.05/8, 4 ROIs and 2 frequency-windows of interest). We also ran this analysis across all frequency-points (4 to 30Hz) however this did not reveal any effects in any ROI.

**Figure 5.**
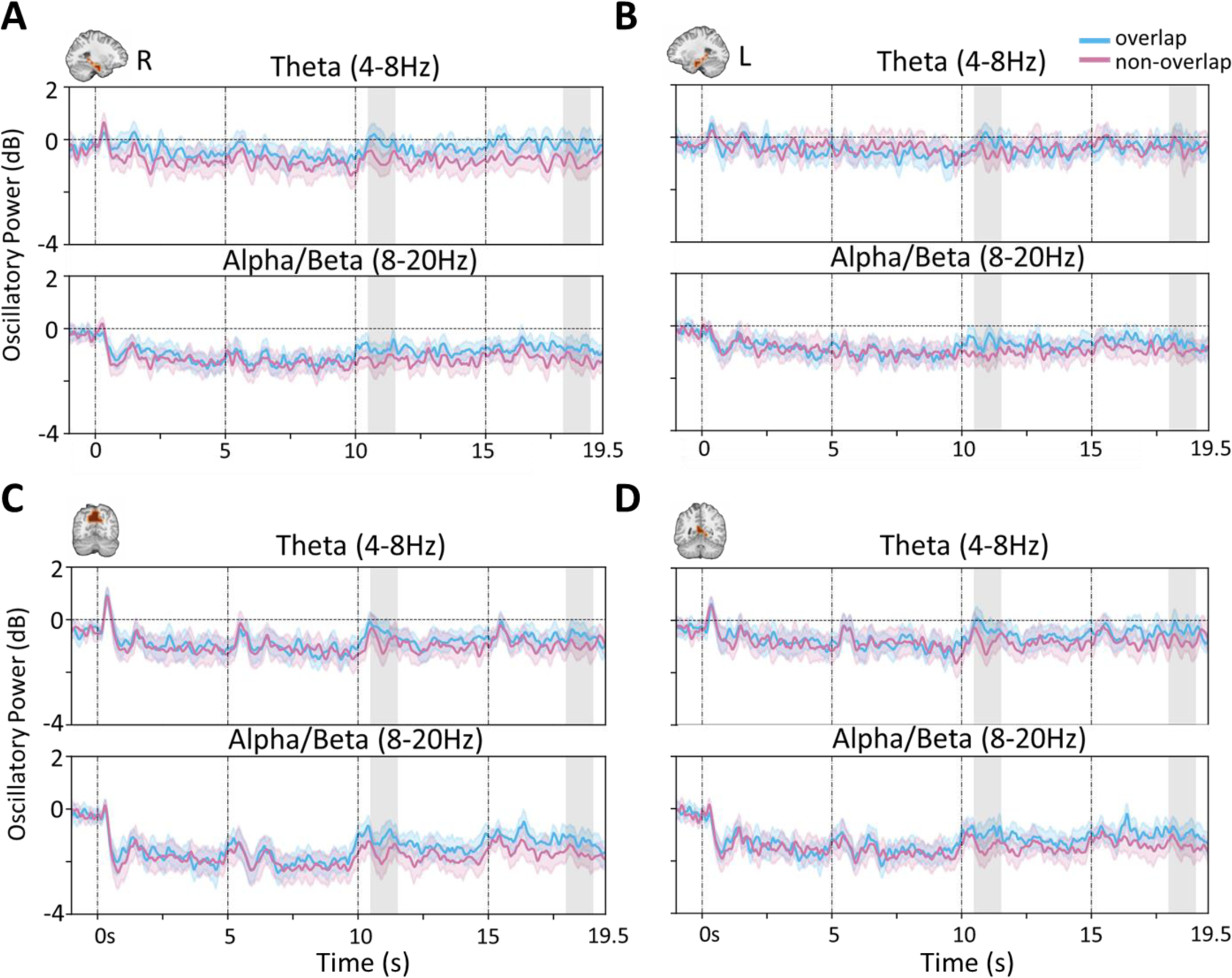
Oscillatory power averaged across theta (top) and alpha/beta (bottom) bands and plotted from video-onset to video-offset (0 to 19.5s) in the right hippocampus (A), left hippocampus (B), precuneus (C) and retrosplenial cortex (D). Grey shaded regions depict 10-11.5s and 18-19.5s post-video onset, where sensor-level effects were found.

In sum, we found that oscillatory power in theta and alpha/beta bands was greater when participants watched overlap videos compared to non-overlap videos, especially during the second repetition of the video (10-20s). These effects were localised to the hippocampus, precuneus, and retrosplenial cortices among other default mode network regions.

#### Subsequent memory effect

Finally, we ran paired-samples t-tests to detect potential differences in oscillatory power between subsequently remembered vs. forgotten locations, as measured in the endpoint-matching task. Oscillatory power was averaged over time-frequency points that belonged to the overlap vs. non-overlap clusters for each ROI (reported above) and compared between subsequently remembered and forgotten locations. We included trials where participants could successfully match both endpoints (A/B) when cued by the other, and trials where they remembered either endpoint B cued by endpoint A or vice versa but were incorrect on the other test (endpoint A cued by endpoint B). Note, we only performed these analyses in the ROIs where significant overlap vs non-overlap differences were seen.

In the 10 to 11.5s time-window, there was a significant effect in the left hippocampus, with subsequently remembered videos showing lower oscillatory power than subsequently forgotten videos (*M_remembered_* = −0.94, *SD* = 1.09; *M_forgotten_* = -.11, *SD* = 1.39; *t*(23) = −2.07, *p* = 0.049, *d* = .43). No significant differences in oscillatory power between subsequently remembered and forgotten locations were seen in the right hippocampus (*M_remembered_* = -.32, *SD* = .78; *M_forgotten_* = -.35, *SD* = 1.46; *t*(23) = .08, *p* = .96) or precuneus (*M_remembered_* = −1.05, *SD* = 1.33; *M_forgotten_* = −1.21, *SD* = 1.62; *t*(23) = .47, *p* = .63). There was no significant effect in the right hippocampus in the 18 to 19.5s window (*M_remembered_* = −1.06, *SD* = 1.49; *M_forgotten_* = -.43, *SD* = 1.21; *t*(23) = 1.47, *p* = .16). None of these effects survive Bonferroni correction for multiple comparisons (p<0.05/4). Therefore, we found that oscillatory power from the theta and alpha bands in the left hippocampus was lower for subsequently remembered compared to subsequently forgotten videos (though we note the smaller sample size of 24 and marginal p-value). The left hippocampus was the only ROI to show a significant subsequent memory effect.

## Discussion

We used a scene-integration paradigm to determine the region-specific neural oscillations involved in forming location-based representations that are critical for navigating new environments. We replicated previous behavioural findings that participants were able to infer which scenes came from the same location following exposure to panoramic videos depicting the spatial relationship between two perceptually distinct viewpoints of the same location. We found changes in neural oscillatory power in the theta (4-8Hz) and alpha and low beta (8-20Hz) frequencies in the overlap condition, where participants were able to infer which two viewpoints were from the same location, relative to the non-overlap condition, where participants were not able make the same location-based inference. These oscillatory power changes were localised to the hippocampus and to other scene-selective regions including the retrosplenial cortex and precuneus. Therefore, we have identified the region-specific oscillatory dynamics involved in participants ability to infer common locations from differing viewpoints.

We were unable to provide evidence, using pattern classification analyses, for changes in neural patterns as a function of watching the videos (indeed, we were not able to use pattern classification analyses to dissociated individual scenes). Thus, unlike in our previous fMRI study (Berens et al., 2021), we were not able to provide more direct evidence for the emergence of ‘location-based representations’. However, given our previous fMRI study provided evidence for these neural representations, and that their emergence was associated with participants ability to infer two viewpoints were from the same location, it is highly likely that such representations underpinned behaviour in the present experiment (given we saw the same behavioural pattern across the two independent experiments). Thus, we continue to refer to ‘location-based representations’ throughout the manuscript, with the caveat that we infer their presence with reference to our previous fMRI study.

MEG data collected when participants watched videos revealed greater theta power for overlap than non-overlap videos. Theta increases were maximal in parietal sensors and were estimated to arise from several regions in the brain including the hippocampus. Therefore, we find evidence that hippocampal theta increases were found when participants viewed videos that allowed them to integrate two perceptually distinct scenes into a location representation based on spatial context. Our findings are in line with a previous study that found hippocampal theta increases using MEG when participants learned a hidden location in a virtual Morris water maze compared to when they randomly moved around in the maze, when no location memories were formed (Pu et al., 2017). Theta increases were also found when participants formed associations in a spatial context compared to a baseline, pre-stimulus period, during fixation cross presentation (Crespo-García et al., 2016). We extend these findings by showing that theta power increases were involved when participants were attempting to integrate two scenes into one coherent location-based representation, in the absence of any overt navigation task. This evidence therefore adds to the literature implicating the hippocampal theta rhythm in memory formation and spatial navigation (Backus et al., 2016; Caplan & Glaholt, 2007; Crespo-García et al., 2016; Griffiths et al., 2016; Joensen et al., 2023; Staudigl & Hanslmayr, 2013; Summerfield & Mangels, 2005).

Critically, this theta power difference occurred during the second repetition of the panoramic video within a trial. Theta power increased around 500ms following the start of the camera pan from the endpoint, and increased as the camera pan approached the central overlapping scene between the two endpoints. The timing of this theta power increase is likely important. First, its presence in the second repetition of the panoramic video suggests that a theta-mediated process is occurring only after participants have been presented with information that allows them to infer the two endpoints are from the same location (in the first repetition of the panorama). This perhaps suggests the theta power differences relate to the formation of a location representation as opposed to the simple awareness of a potential relationship. An online assessment of this awareness, tracking when participants become fully aware that the video is depicting a coherent location, would allow us to assess the timing of theta power increases relative to this awareness in future research.

Second, theta power changes occurred when participants saw the endpoints of the panorama. One possibility is that the viewpoint not currently presented is being actively retrieved and associated/integrated with the currently presented viewpoint and theta is tracking that retrieval and integration process. This would fit with evidence that theta is associated with not just associative memory (i.e., encoding A-B pairs presented at the same time; Caplan & Glaholt, 2007; Hanslmayr & Staudigl, 2014; Staudigl & Hanslmayr, 2013; Summerfield & Mangels, 2005) but also associative inference where participants integrate separately encoded A-B and A-C pairs (Backus et al., 2016). This integration process is thought to be underpinned by the retrieval of non-presented but associated information (i.e., retrieving B when learning A-C), resulting in the integration of the presented and retrieved information (Molitor et al., 2021; Morton et al., 2023). A similar process could be occurring when participants view each endpoint, but only once they realise the endpoints are from the same location. Using pattern classification to assess the reactivation of endpoints during video presentation could assess this possibility. We were unable to distinguish between scene endpoints in the present study, preventing us from running this analysis. Future research would likely need fewer scenes with more repetitions per scene and more repetitions per video to allow for this type of analysis.

The vMPFC is also considered to interact with the hippocampus during this inferential integration process (Morton et al., 2023; Preston & Eichenbaum, 2013), possibly via the theta rhythm (Backus et al., 2016). We did find that increases in theta power for overlap videos compared to non-overlap videos were localised to the vMPFC, but the vMPFC – Hippocampus connectivity analyses proved inconclusive, possibly due to low statistical power (see Supporting Information for details regarding the connectivity analyses). Theta power increases that we found in the vMPFC and the hippocampus could thus reflect similar inferential learning between the two endpoints to form location-based representations.

Another possibility is that both endpoints A and B are integrated into a more abstract, allocentric representation of the location (A*) that is reactivated during the memory test and leads to successful matching of endpoints A and B (Berens et al., 2021). Previous theories and computational models predict that such spatial learning occurs via communication between medial parietal and temporal regions in the scene-selective network (Baumann & Mattingley, 2021; Bicanski & Burgess, 2018; Byrne et al., 2007; Marchette et al., 2014). In line with these theories, we find that changes in oscillatory power during the formation of location-based representations were localised to the hippocampus, retrosplenial cortex, precuneus, lateral parietal cortices (IPL and SPL) and early visual cortex. Previous studies using similar paradigms have used fMRI and have found that the retrosplenial and parahippocampal cortices are responsible for representing location-based information (Berens et al., 2021; Robertson et al., 2016). However, these studies were not able to conclude that these regions were involved in the formation of these representations. It is not clear whether participants formed a more abstract allocentric representation (i.e., A*), or whether they formed a simpler associative memory (i.e., A-B), or whether the integrative mechanisms discussed above are the mechanistic basis for forming either of these possible representations. However, our results point clearly to a role for hippocampal and medial parietal theta in forming a more location-based representation that can support behaviour in the scene-matching task.

We also investigated the neural oscillations involved in the successful formation of location-based representations by contrasting activity from overlap videos that were subsequently remembered vs. forgotten (Subsequent Memory Effect, SME). We found hippocampal theta decreases for remembered compared to forgotten locations, hence we found a negative SME. This effect was just below the significance threshold (*α* = .05), possibly due to low trial and participant numbers leading to a low signal-to-noise ratio and thus low statistical power to detect effects. As such, compared to our main analyses that are powered appropriately, we draw caution in relation to this result (i.e., this result could simply be a false positive).

Several studies have reported increases in theta power for subsequently remembered compared to subsequently forgotten stimuli, for both item (Fellner et al., 2013; Guderian et al., 2009; Hanslmayr et al., 2009, 2011; Khader et al., 2010; Meeuwissen et al., 2011; Osipova et al., 2006; Sederberg et al., 2006) and associative memory (Backus et al., 2016; Caplan & Glaholt, 2007; Crespo-García et al., 2016; Griffiths et al., 2016; Joensen et al., 2023; Staudigl & Hanslmayr, 2013; Summerfield & Mangels, 2005). However, studies that have investigated associative learning in a spatial context have observed decreases in theta power for subsequently remembered vs. forgotten items. For example, Fellner et al., (2013) conducted a study where participants memorised a list of items using the method of loci, which involves associating items together in a spatial context, vs. the pegword method, which involves associating items together in a non-spatial context of a rhyme. They found a negative subsequent memory effect for items associated together in a spatial context compared to a non-spatial context. Two other studies on theta changes during associative learning in a spatial context also found theta decreases (Crespo-García et al., 2016; Griffiths et al., 2016). They studied item and location binding as participants navigated around a specified path and found theta decreases for subsequently remembered locations, cued by the items, compared to subsequently forgotten locations. Therefore, it is possible that negative subsequent memory effects arise when encoding of information occurs in a spatial context.

If a greater decrease in theta does reflect better encoding (and therefore subsequent memory), it is unclear why we also saw even greater decreases in theta in the non-overlap condition, where no learning appeared to be occurring (i.e., participants couldn’t match the scenes presented in the non-overlap condition following video watching). Although speculative, one possibility is that the decrease in theta we observe reflects a form of ‘encoding effort’. In the overlap condition, more ‘effort’ results in better subsequent memory as a location-based representation can be formed. However, in the non-overlap condition, increased ‘effort’ does not result in the formation of a location-based representations (as the scenes are from different locations) resulting in poor behavioural performance.

As well as seeing theta differences between the overlap and non-overlap condition (and a theta SME), we also found that alpha and low beta power (12 to 20Hz) was greater during the overlap than non-overlap condition. A recent study found that alpha/beta power increased during active environmental encoding compared to guided encoding and attributed this increase to increased attention during active vs. passive exploration (Chrastil et al., 2022). Therefore, it is possible that participants paid more attention while watching overlap than non-overlap videos. However, the dominant finding in the literature is that stimulus-onset induces a decrease in alpha/beta power that is considered to reflect increased attention (Klimesch, 2012) and representing stimulus-information necessary for the formation of episodic memories (Griffiths et al., 2021, 2019; Hanslmayr et al., 2016; Parish et al., 2018). Thus, increased alpha/beta in the overlap condition is unlikely to be driven by simple attentional differences, unless participants were paying greater attention to the non-overlap condition (perhaps because it is more difficult to associate two scenes from different locations). These alpha/beta changes source-localised to the same hippocampal and medial parietal regions that were revealed in the theta analyses, although the precuneus perhaps was driven more by alpha/beta than theta changes suggesting a potential dissociation.

Our results are also in line with previous theories implicating the hippocampus, retrosplenial cortex and precuneus as critical for maintaining and transforming location-based representations (Alexander et al., 2023; Bicanski & Burgess, 2018; Parish et al., 2018). This model proposes that ego-centric representations are supported by the precuneus and allocentric representations are supported by the hippocampus, with the retrosplenial cortex acting as a transformation circuit between these two references frames. There is evidence for more allocentric place cells in the hippocampus (O’Keefe & Dostrovsky, 1971) and a heterogeneity of neural representations in the retrosplenial cortex, including neurons that fire in relation to combinations of heading direction, location, and landmark information (Chen et al., 1994; Knight et al., 2014; Lozano et al., 2017; Mao et al., 2017). Other theories implicate the hippocampus in allocentric ‘scene construction’ that supports both spatial navigation and episodic memory (Hassabis & Maguire, 2007) and the retrosplenial cortex in landmark learning (e.g, Auger et al., 2012, 2015; Auger & Maguire, 2018) and the learning of schematic spatial knowledge (Farzanfar et al., 2022; Mitchell et al., 2018; Peer & Epstein, 2021). Despite their differences, all these theories implicate the hippocampus and retrosplenial cortex in the learning of spatial representations.

We found oscillatory changes during location-learning in these brain regions, broadly in support of these theories. We did not however see clear regional differences in relation to oscillatory frequency bands, nor did we track the information flow across these regions over time. Such analyses would provide more mechanistic evidence for these theories. Future research could focus on hippocampal theta vs. medial parietal alpha/beta in relation to how these regions communicate within these frequency bands during the learning of novel spatial environments (Hanslmayr et al., 2016; Parish et al., 2018). Additionally, future research could investigate how these regions communicate via oscillations using functional or effective connectivity measures to provide more concrete evidence for these theories.

Although our effects were significant after correcting for multiple comparisons within each time-window, some of the effects did not survive a more stringent Bonferroni correction across time-windows. Specifically, our sensor-level effects failed to survive Bonferroni correction across the 6 time-windows of interest, and the non-hippocampal source-level effects also did not survive this correction. We did find that the hippocampal source-level results are robust since they are present in every analysis that we conducted. Therefore, we interpret the theta/alpha oscillatory changes in the hippocampus during the formation of location memories as a robust effect. It is important to note that the Bonferonni correction is especially conservative in this analysis because the data across the time-windows are likely not entirely independent. Figure 5 suggests the possibility of a relatively sustained (but small) increase in theta/alpha from 10-20secs. As such, even though our time-windows are not temporally adjacent, we may be assessing oscillatory differences that are driven by the same underlying source. This is also the first study to investigate the neural oscillations related to the formation of location-based representations. Considering these two points, we present the effects that survive correction within each time-window but have made clear which of these effects do not survive more stringent corrections across time-windows.

In conclusion, participants were able to match perceptually distinct viewpoints from the same location after viewing videos that showed the spatial relationship between these endpoints, potentially driven by the formation of a coherent representation of that location. Using MEG, we found that hippocampal and medial parietal theta and alpha/beta power increased when participants watched these videos compared to videos that did not allow for location-based representations to be formed. This finding adds to the literature implicating the hippocampal theta rhythm in episodic memory formation and spatial navigation (Buzsáki & Moser, 2013; Nyhus & Curran, 2010) by showing that it is implicated in combining egocentric, viewpoint-dependent scene information into a potentially allocentric, viewpoint-independent representation. Thus, we provide evidence in favour of models that predict the involvement of hippocampal and medial parietal regions in the formation of location-based representations (Bicanski & Burgess, 2018; Byrne et al., 2007), and implicate theta and alpha/beta oscillations in the formation of such representations.

## Acknowledgements

This work was funded by a Wellcome Trust Institutional Strategic Support Fund (ISSF) to The University of York (204829/Z/16/Z), through the Centre for Future Health (CFH). AJH is funded by the Economic and Social Research Council (ESRC; ES/X00791X/1) and The Leverhulme Trust (RPG-2023-003). The authors thank Benjamin Griffiths, Richard Aveyard, Dan Denis, and Pranay Yadav for their advice with data analysis and the York Neuroimaging Centre (YNiC) staff for their assistance during data collection. For the purposes of open access, the authors have applied a creative commons attribution (CC-BY) license to any author accepted manuscripts version arising henceforth.

## Data availability statement

The behavioural data and analysis code, and MEG analysis code that support the findings of this study are openly available in the Open Science Framework [here](https://osf.io/y4hev/?view_only=fce12d0877d4402ca54d61923ca3bf9e).

