## Supplemental Information for "Theta and alpha oscillations in human hippocampus and medial parietal cortex support the formation of location-based representations"

### GLMM Methods and Results

Behavioural data from the endpoint matching tasks were also analysed by fitting generalised linear mixed-effects models (GLMMs) using the glmer software from the lme4 package (Bates et al., 2014) implemented in R. We analysed the data in this way to most closely replicate the analysis in Berens et al., (2021). This modelling approach makes fewer statistical assumptions than traditional analyses of variance (ANOVAs) and can deal more robustly with non-normal and non-independent data (Baayen et al., 2008). Moreover, by-item and by-participant variance can be estimated in a single statistical model, while also being more appropriate for the analysis of categorical data (such as correct [1] vs. incorrect [0] in the present experiment) than ANOVAs (Jaeger, 2008). We therefore present these analyses to complement the more standard analyses presented in the manuscript.

For the endpoint matching task, the fixed effects (i.e., predictors of interest) were Session (1: pre-videos, 2: post-videos) and Condition (non-overlap, overlap, unseen). Both factors were dummy-coded, with session 1 and non-overlap as the reference categories. Random effects were added for Participants and Items (endpoint images).

The GLMMs were built by starting out with the maximal random effects structure justified by the design of the experiment (Barr et al., 2013), before systematically reducing the random effects structure to increase the power of the analysis (Matuschek et al., 2017). This was achieved by removing random slopes/intercepts and continually comparing the new (reduced) model to the former (full) model, aiming for the simplest model (in terms of its random effects structure) that is not statistically different from the maximal model regarding how well it fits the data. Model comparisons were made using the *anova* function (testing whether a reduction in the residual sum of squares is statistically significant), with a chisq-threshold of < 3. For all GLMMs, the "bobyqa" optimiser was used to increase the likelihood of model convergence. After choosing the best-fitting model, the *emmeas* package was used to conduct pairwise comparisons on the GLMM to aid the interpretation of significant interactions (adjusted for multiple comparisons with Holm correction). The analysis code is available on OSF.

GLMMs were fit to participants' trial-by-trial responses during the endpoint-matching task. The GLMM that explained the most variance in our data included random intercepts for both participants and items. Models with more complex random-effects structures (e.g., including random slopes) either resulted in singular fit issues or did not fit the data as well as the simpler random-intercepts-only model. The main effects were non-significant (*zs* < .86, *ps* > .39). Importantly, a significant interaction between post-video watching and Condition was found, $z = 2.52$, $p = .011$. Follow up paired tests revealed that participants were able to match endpoints more successfully post- vs. pre-videos in the overlap condition ($z = 4.55$, $p < .0001$) but not non-overlap ($z = 0.85$, $p = .65$) or unseen conditions ($z = 0.98$, $p = .65$). Therefore, participants learnt locations better after watching overlap videos compared to non-overlap videos and unseen locations. These results match the main analyses.

### Phase-based functional connectivity

Previous research indicates that the ventromedial prefrontal cortex (vMPFC) and hippocampus communicate via the theta rhythm during memory integration and spatial imagery (Backus et al., 2016; Colgin, 2011; Kaplan et al., 2017; Preston & Eichenbaum, 2013). Therefore, we tested for functional connectivity between the mPFC and hippocampus during the formation of location-based representations in this paradigm. We also expected the hippocampus to interact with the medial parietal regions as part of the scene-selective network (Baumann & Mattingley, 2021; Epstein, 2008; Marchette et al., 2014; Vass & Epstein, 2013).

We tested these predictions by running a seed-based phase synchronisation analysis in the following way: First, the signal at each voxel in the brain was band-pass filtered between 4 to 8Hz (theta band) and the instantaneous phase was derived at each time-point, at each voxel, by computing the Hilbert transform signal. A voxel in the mPFC (for hippocampal <-> mPFC connectivity, (Backus et al., 2016; Kaplan et al., 2017) and in the hippocampus (for hippocampal <-> medial parietal lobe connectivity) region-of-interest with the greatest difference in oscillatory power between overlap and non-overlap conditions was chosen as the seed. Second, the phase-lag index (PLI) was then calculated between that seed voxel and every other voxel in the brain, resulting in a map of PLIs in each voxel of the brain. The absolute PLI value ranges between 0 to 1 with values closer to 1 indicating high connectivity and 0 indicating no connectivity. Cluster-based permutation tests were then used to detect voxels with statistically significant differences in PLI 1) between overlap and non-overlap video watching during the time-windows, and 2) greater than 0 in the overlap condition.

No clusters were statistically significant after correcting for multiple comparisons. The largest cluster that was above the (uncorrected) significant threshold ($p$ = .103 corrected p-value; for the mPFC seed) included the SMA and pre-SMA, and posterior-cingulate cortex including the retrosplenial cortex.

**Multivariate analysis of Oddball Detection Task**

We conducted multivariate analysis of sensor-level MEG data recorded as participants were passively viewing the endpoints with two aims: 1) to replicate Berens et al.'s (2021) findings that pattern similarity between two endpoints from the same location increases after, relative to before, watching overlap videos, and 2) to test for potential reactivation of the other endpoint from the same location after perceiving an endpoint in the video, based on previous studies that have found such reactivation of to-be-integrated information during episodic memory formation (Molitor et al., 2021; Morton et al., 2023). We used representational similarity analyses (RSA) to test hypothesis 1 and trained pattern classifiers to test hypothesis 2.

Before testing the theoretical hypotheses, we aimed to determine if either RSA or a pattern classifier could distinguish between individual endpoints. Neither of these methods could differentiate patterns of activity from the same endpoint relative to other endpoints. Therefore, we did not test our theoretical hypotheses further. We believe that these analyses failed due to low trial numbers per endpoint. Nevertheless, the analysis steps are described below.

Representational similarity analyses were conducted for each red dot task separately, with the following steps: first, the trials for each endpoint were split into two sets (A and B) using random sampling (without replacement). Then, a grand-average waveform was calculated for each set and each endpoint, resulting in one grand-averaged waveform for set A and set B, for each endpoint. Representational similarity analysis was then conducted by correlating the sensor topology between two grand-averaged waveforms at each time-point of the waveform resulting in a time-series waveform of similarity values. We derived two types of similarity waveforms: 1) Within-endpoint similarity, calculated by correlating the waveforms derived by the two sets of trials from the same endpoint; and 2) Between-endpoint similarity, calculated by correlating two sets of trials from different endpoints. If there is unique information about a specific endpoint in the patterns of MEG activity, then the similarity between two sets of trials from the same stimulus should be greater than the similarity between two sets of trials from different stimuli. However, we did not find any statistically significant difference between within-endpoint and between-endpoint similarity waveforms. Hence, we could not isolate the representation of a unique endpoint in our dataset with this analysis technique.

We also attempted to train a classifier (logistic regression with lasso regularisation) on the data from the Red Dot task in the following way: we pooled trials for each endpoint across pre- and post-video watching to improve signal-to-noise ratio. The data were then downsampled to 100Hz. The classifier was trained on data from 100ms to 300ms with all 244 MEG sensors as features. The classifier was trained and tested on separate sets of trials using k-fold cross validation (6 folds were used in this analysis). However, the classifier could not accurately predict the identity of an endpoint, with accuracy near chance level. It is likely that that these analyses failed due to low signal-to-noise ratio because of insufficient trial numbers. Future research should have more repetitions of fewer endpoints to increase power.
